# Clonal evolution during metastatic spread in high-risk neuroblastoma

**DOI:** 10.1101/2022.08.15.503973

**Authors:** Gunes Gundem, Max F. Levine, Stephen S. Roberts, Irene Y Cheung, Juan S. Medina-Martínez, Yi Feng, Juan E. Arango-Ossa, Loic Chadoutaud, Mathieu Rita, Georgios Asimomitis, Joe Zhou, Daoqi You, Nancy Bouvier, Barbara Spitzer, David B. Solit, Filemon Cruz Dela, Michael P. LaQuaglia, Brian H. Kushner, Shakeel Modak, Neerav Shukla, Christine A. Iacobuzio-Donahue, Andrew L. Kung, Nai-Kong V. Cheung, Elli Papaemmanuil

## Abstract

High-risk neuroblastoma is generally metastatic and often lethal. Using genomic profiling of 470 sequential and spatially separated samples from 283 patients, we characterize subtype-specific genetic evolutionary trajectories from diagnosis, through progression and end-stage metastatic disease. Clonal tracing timed disease initiation to embryogenesis. Continuous acquisition of structural variants at disease defining loci (*MYCN, TERT, MDM2-CDK4*) followed by convergent evolution of mutations targeting shared pathways emerged as the predominant feature of progression. At diagnosis metastatic clones were already established at distant sites where they could stay dormant, only to cause relapses years later and spread via metastasis-to-metastasis and polyclonal seeding after therapy.

## Introduction

Neuroblastoma is an embryonal tumor arising from the developing sympathetic nervous system accounting for 15% of pediatric cancer mortality^1^. Disease presentation is highly heterogeneous and ranges from low-risk local-regional tumors to widely disseminated high-risk disease seen in two thirds of the patients. For high-risk neuroblastoma (HR-NB), modern clinical management includes multimodal chemotherapy, surgical resection, radiotherapy and immunotherapy. Nevertheless, despite intensive treatment 50% of HR-NB patients still relapse with fatal outcomes^2^.

Notwithstanding the metastatic nature of HR-NB, most genomic studies of disease progression focused on small cohorts of paired diagnostic-relapse tumors^3–9^. Recently, broad copy number aberrations (CNA) and whole-exome sequencing (WES) from multi-region biopsies suggested elevated genetic heterogeneity in high-risk disease^10–12^. However, the majority of disease-defining alterations in HR-NB^13,14^ result from structural variants (SVs) that cannot be captured by low-resolution WES/CNA analysis. Todate, the temporal and spatial genomic features of disease progression as patients go through multiple lines of therapy are not well understood. Here, we leverage the MSKCC neuroblastoma biobank to study a unique cohort of 470 tumors from 283 patients representative of HR-NB at diagnosis, consecutive relapses, and diverse metastatic sites. Using a combination of whole genome (WGS) and targeted sequencing approaches we characterized the composite genetic alterations associated with neuroblastoma pathogenesis and define the lineage relationships during disease progression across diverse neuroblastoma molecular subtypes.

## Results

### Cohort ascertainment

Our cohort consisted of 470 tumors from 283 patients with predominantly stage-4 disease (87%) and/or spatially and temporally separate tumors available (Fig. 1a and Supplementary Table 1). Fresh-frozen surgical specimens were collected at consecutive clinical intervention time points including: 110 pre-treatment diagnostic samples, 5 therapy-naive re-resections, 132 therapy resection during induction chemotherapy (t-resection), 111 relapses and 112 further relapses (1-17 samples per patient) and spanned spatially separated disease sites including: 217 samples from primary site, 150 from local-regional spread and 97 from metastases as defined by clinical guidelines^15^ including rare metastatic sites such brain. A web portal describing the treatment timelines and clonal phylogenies for patients with two or more tumors is provided in (https://master.d32nckcows37aj.amplifyapp.com/) and detailed summaries of genomic and clonal evolution patterns in 45 patients with multi-WGS data are provided in Supplementary Information.

**Figure 1.**
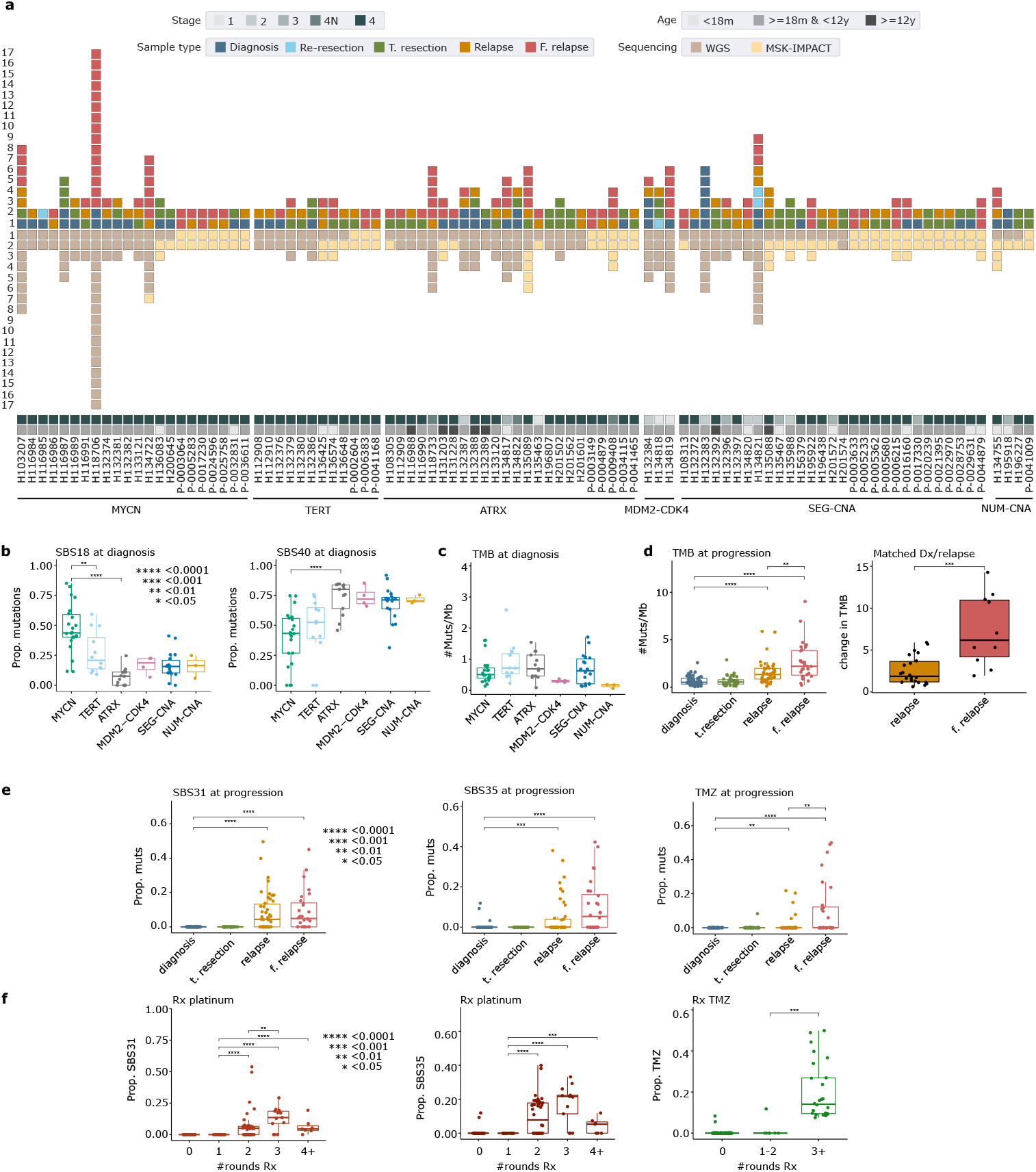
Patient cohort and genome-wide mutational landscape. **a)** Barplot shows the number of tumors sequenced from 94 patients with two or more samples color-coded by type of sample and sequencing performed. Patients are shown as columns organized by the disease subtype which are as follows: 1) MCYN, patients with *MYCN* amplification. 2) TERT, patients with *TERT* SVs. 3) ATRX, patients with *ATRX* events. 4) MDM2-CDK4, patients with *MDM2-CDK4* co-amplifications. 5) SEG-CNA and 6) CUM-CNA, patients with segmental or numeric CNAs but without the aforementioned alterations. (SV, structural variants. CNA, copy number aberration. WGS, whole genome sequencing.). Information about age and stage at diagnosis is provided as tile plots at the bottom. Box plots show comparison of the proportion of mutations attributed to SBS18 and SBS40 **(b)** and tumor mutation burden (TMB) **(c)** in WGS data across diagnostic tumors of different molecular subtypes (n=72 tumors). **d)** Box plot on the left shows the increase in TMB across samples collected at diagnosis, t-resection, relapse and further relapse (n=129 patients) while on the right is the fold change in TMB in relapse and further relapse samples compared to the matched diagnostic tumor of the same patient (n=22 patients). **e)** Box plots show the proportion of substitutions attributed to SBS31, SBS35 and temozolomide (TMZ) signatures across samples collected at diagnosis, t-resection, relapse and further relapse (n=129 patients). **f)** Boxplots show the number of substitutions attributed to therapy-related mutational signatures for tumors from patients with stage-4 disease who were exposed to increasing numbers of rounds of platinum or temozolomide-based chemotherapy (n=145 tumors). The data and script for Fig. 1 are available in Supplementary Table 1 and the GitHub repository.

### Landscape of genomic alterations

Comprehensive genomic profiling (substitutions, indels, SVs and CNAs) (Extended Data Fig. 1a-b, Supplementary Table 2) identified both established and novel gene mutations linked to neuroblastoma pathogenesis. The genomic landscape was representative of HR-NB^16,13,17^. Only 4 genes (*MYCN, ALK, TERT, ATRX*) had mutations in >10% of patients while recurrent *MDM2-CDK4* co-amplification was observed in 3%^18^. Alterations in *MYCN, TERT, ATRX* and *MDM2-CDK4* defined mostly non-overlapping disease subtypes explaining 51% of the cohort while *ALK* mutations were shared across the cohort. 41% and 8% of the patients did not have any subtype-defining alterations other than segmental (SEG-CNA) or numeric chromosome-level CNAs (NUM-CNA), respectively.

*TERT* rearrangements are common in neuroblastoma^13,14^. Here, we report *TERTp* substitutions in 9 patients comprising 17% of the *TERT* events. However, contrary to *TERT* SVs which are mutually exclusive to other subtype defining events, *TERT*p mutations were enriched in *MYCN*-amplified (*MYCN*-A) neuroblastoma and demarcated a group of *MYCN*-A patients with a trend for rapid progression and death within 2 years from diagnosis (Extended Data Fig. 1c). PI3K-mTOR pathway was also mutated in 5% of the patients suggesting enrichment in HR-NB compared to primary neuroblastoma (<1% in Brady et al^14^). Additionally, mutations in neuroblastoma differentiation genes including *RARA, RARB, PHOX2B, SPRY2, IGF2BP3* and *WNT5A* were observed in 4% of the patients at relapse (Supplementary Information).

### Evolution of mutational landscape in response to therapy

Analysis of genome-wide mutation landscapes revealed distinct mutational patterns at diagnosis and relapse (Extended Data Fig. 2a-b and Supplementary Table 1). At diagnosis, two substitution signatures, SBS40 and SBS18, were differentially enriched across molecular subtypes. Mutations attributed to SBS40, similar to the clock-like signature SBS5^19^, was higher in *ATRX*-mutated patients and correlated with age at diagnosis, while SBS18, which is predominantly defined by C>A mutations, prevailed in *MYCN*-A tumors in agreement with prior literature^14^ and did not correlate with age (Fig. 1b and Extended Data Fig. 3a). SBS18 was first described in neuroblastoma^20^ and attributed to reactive oxygen species (ROS)^21^. *MYCN*-A enhances glutaminolysis in neural crest progenitor cells, which in turn induces oxidative stress by ROS production^22^. Expression of glutaminolysis signature was higher in *MYCN*-A tumors compared to other subtypes (Extended Data Fig. 3b). This provides a plausible mechanistic link between *MYCN*-A, metabolic reprogramming and SBS18 burden. Notably, the glutaminolysis gene expression signature and the rate of accumulation of SBS18 remained stable during disease progression (Extended Data Fig. 3b).

Pediatric tumors are defined by low tumor mutation burden (TMB)^23^. TMB was low at diagnosis^24,23^ (median=0.52 muts/Mb, range=0.06-2.6) (Fig. 1c) but increased significantly during disease progression (median=2.2 muts/Mb, range=0.2-9). Notably, in patients with matched diagnostic/relapse tumors, TMB increased by 6- and 14-fold at first and later relapses (Fig. 1d), respectively, approximating TMB ranges seen in adult tumors^20^. At diagnosis neuroblastoma is characterized by an immune-cold tumor microenvironment^25–27^ (TME) with poor responses to immune checkpoint blockade therapy^28,29^. We evaluated whether the increased TMB during disease progression presented a therapeutic opportunity mediated by putative neo-antigens^30^. However, despite the increase in predicted neoantigen burden, there was no association with transcriptional patterns suggestive of an immunomodulatory switch during disease progression (Extended Data Fig. 3c).

At relapse, increase in TMB was associated with exposure to chemotherapy-associated mutation signatures^31^. Specifically, three substitution signatures (TMZ, SBS31 and SBS35) dominated by T>C and C>T mutations correlated with exposure to temozolomide and platinum with evidence of strong dose-response relationships^32,21,31,19^ (Fig. 1e-f). At disease progression, tumors from prior radiated sites had an excess of small deletions, SV deletions, reciprocal translocations and complex events^33,34^ (Supplementary Fig. 1). This demonstrates that therapy directly molds the mutation landscape of HR-NB tumors. Thus, we next evaluated the effect of these mutation processes in the driver landscape at diagnosis and during disease progression. Of 82 oncogenic substitutions from WGS data, 48 were present at diagnosis and 34 emerged at relapse. Notably, only 12% of the oncogenic substitutions were assigned to a therapy-related signature compared to all relapse-specific SNVs (34%) (Extended Data Fig. 3d).

### Timing the emergence of the initial neuroblastoma clone

For each patient, clonal reconstruction of tumor phylogenies delineated the trunk marked by the mutations found in 100% of malignant cells in all tumors of a patient as well as subclonal events (not on trunk). Trunk represents the most recent common ancestor (MRCA) (Online Methods and Supplementary Information, Supplementary Fig. 2-46 and Supplementary Tables 3-4). The number of clock-like mutations on the trunk can be used to estimate the time of MRCA emergence^35–37^. Across 39 evaluable patients the number of truncal substitutions (trunk length) was low (median=753, range=11-5801) and correlated with age at diagnosis only when disease subtype, stage, and number of tumors were taken into account (Fig. 2a and Supplementary Fig. 47). Low-stage disease tended to have shorter trunks (Fig. 2b) with lengths comparable to the number of substitutions detected in non-malignant clones in bulk placenta also enriched for SBS18 mutations^38^. This suggests that the first malignant clone emerges in similar time frames during embryogenesis. Indeed, chronological timing of MRCA emergence using the clock-like SBS40 mutations^37^ confidently pinpointed an embryonic and post-natal origin in 6 and 8 cases, respectively. For the remaining 25 patients the confidence intervals were large, owing to low numbers of mutations on the trunk (Fig. 2c). Notably, MYCN-A was common in embryonic origin (4/6) while ATRX-mutant disease was enriched for post-natal onset (7/8) especially in patients with *ATRX* truncating mutations (Fig. 2c).

**Figure 2.**
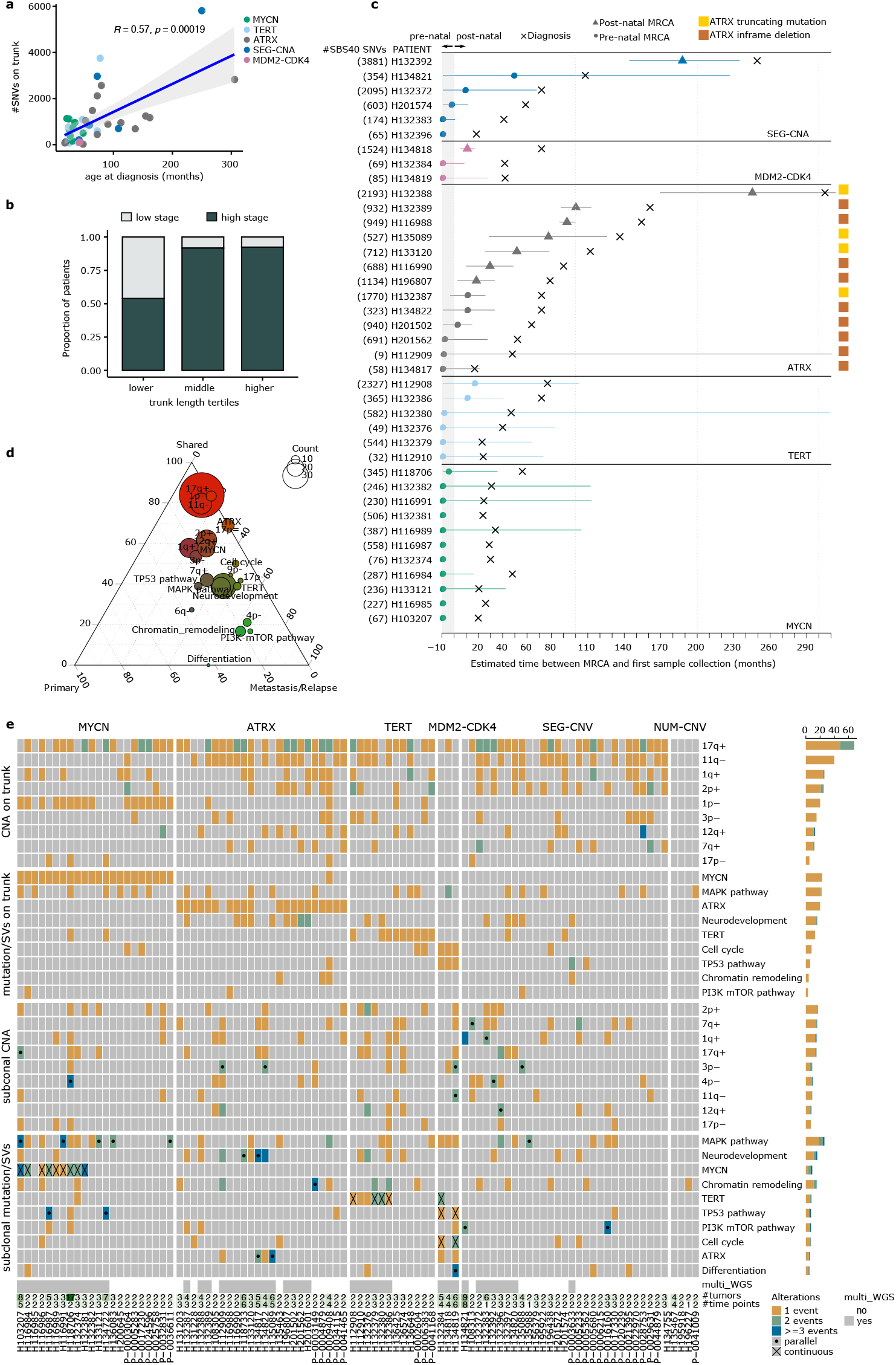
Timing of emergence of the first malignant clone. **a)** Scatter plot shows the relationship between the number of single nucleotide variants (SNVs) on the trunk (trunk length) and age at diagnosis (Pearson correlation). **b)** Barplots show the breakdown of upper, middle and lower tertiles of the trunk length distribution by stage at diagnosis. **c)** Timeline plot shows the time of diagnosis and the predicted time of emergence of the most recent common ancestor (MRCA) with 95% confidence intervals for n=39 patients. Number of SBS40-associated SNVs on the trunk is shown next to the patient id. Shaded area from −9 to 0 on the x-axis shows the period in utero. The predicted time of emergence of the MRCA is shown with a circle (pre-natal) or a triangle (post-natal). **d)** Ternary plot shows the proportion of events that are shared by all tumors of a patient, seen in a subclone specific to a sample from the primary site or metastatic/relapse site. Dots are color-coded red or green to indicate a tendency to be shared or metastasis/relapse-specific with a size proportional to the total number of events. **e)** Heatmap shows the number of truncal and subclonal genetic changes identified in 94 patients with >=2 tumors. Mutations and SVs are collapsed to the affected pathways except for those hitting the most recurrent disease-defining genes (*MYCN, TERT* and *ATRX*). Black dots indicate parallel evolution while crosses indicate the loci affected by continuous subclonal SVs when there is already an SV event on the trunk of the patient. Lowermost tile plot shows the availability of multi-WGS data, number of tumors and the timepoints studied for each patient. Barplot on the left shows the frequency of the events per row. +, gain. -, loss. The data and script for Fig. 2 are available in Supplementary Tables 1, 2 and 4 and the GitHub repository.

### Subtype-specific evolutionary trajectories underwrite disease progression

Analyses of multiple samples representative of the clinical course of treatment from 94 patients across HR-NB subgroups identified previously unappreciated and subtype-specific evolutionary trajectories for tumors with *MYCN*-A, *TERT*-SV, *ATRX* events and *MDM2-CDK4* co-amplification (Fig. 2d-e, Extended Data Fig. 4-5 and Supplementary Data fig. 2-46 and 48-53). While subclonal events at RAS-MAPK^4,5,39^ and PI3K-mTOR pathways were common across subtypes, the acquisition of SVs emerged as critical events in disease evolution with a striking propensity to repeatedly target the main subtype-defining driver genes *MYCN, TERT* and *ATRX* (Fig. 2d-e).

Tumor-initiating *MYCN* amplifications in neuroblastoma^40^ are frequently found in extrachromosomal DNA^41,42^, which may result from simple or complex SVs at the trunk of the tumor phylogeny. Amongst patients with multi-WGS data, we observe continuous rearrangements of the *MYCN* locus in both primary (7/10 cases) and relapse sites (9/11 patients) (Fig. 2e, Fig. 3a and Supplementary Fig. 54-56). During therapy, rearrangements at the *MYCN* locus continued to accumulate in 9/10 patients and in four of these patients a clone with chromothripsis at the *MYCN* locus dominated across metastatic sites without evidence of further diversification (Supplementary Fig. 54-56).

**Figure 3.**
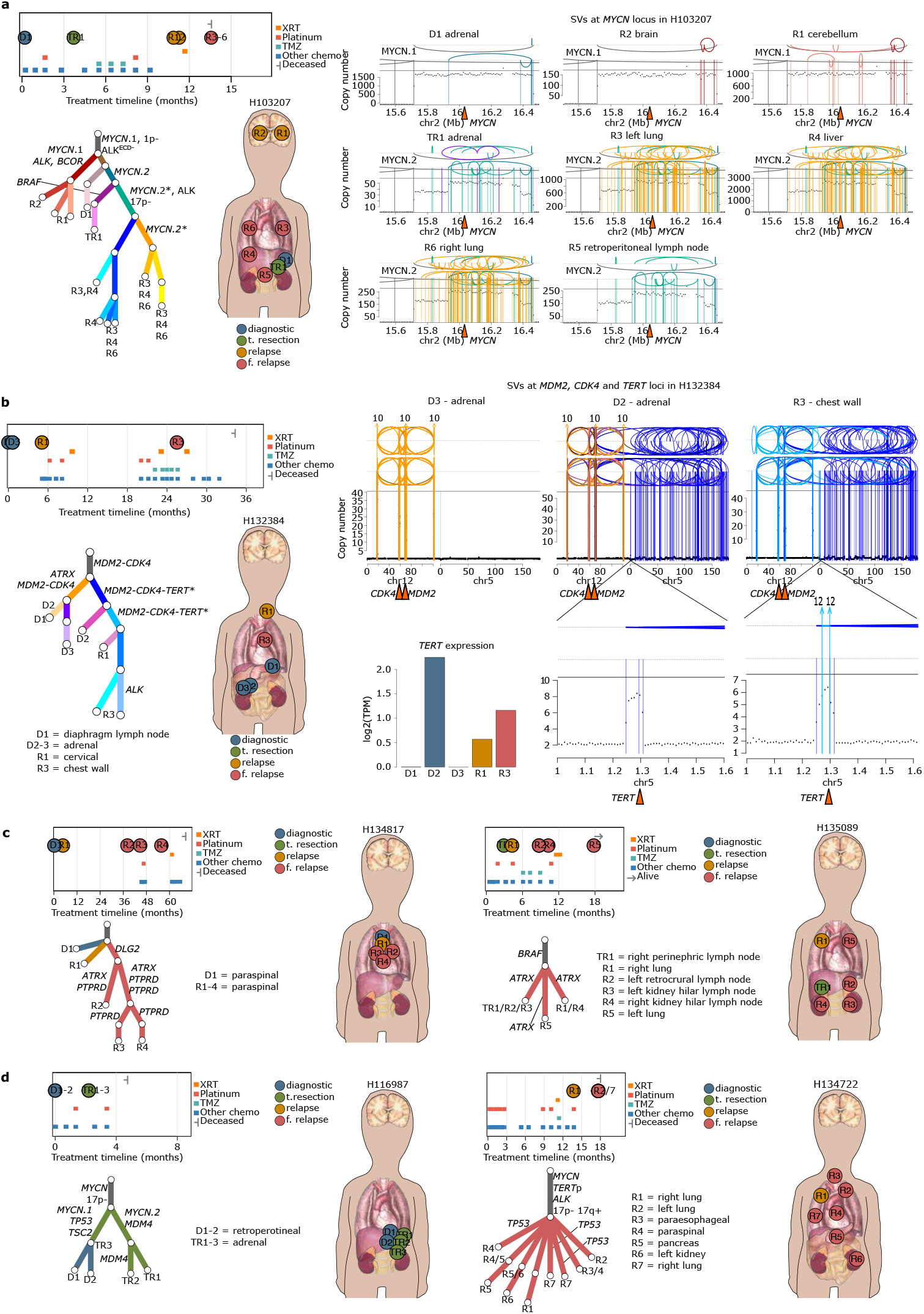
Subtype-specific evolutionary trajectories. **a)** Subclonal structure for patient H103207 is summarized in multiple panels. Treatment timeline gives a summary of the therapy administered, the sequenced tumors and the survival status at last followup. Body map shows the location of the tumor sites sequenced. Subclone tree shows the lineage relationships amongst the subclones identified in a patient. Subclones are designated by branches with non-informative lengths. Trunk is shown in gray at the top of the tree. Terminal nodes are annotated with tumors where the corresponding clone is present. Branches are annotated with putative oncogenic events. Different types of *MYCN* amplicons are indicated by a number after gene name (i.e. *MYCN*.1). Continuous accumulation of SVs at *MYCN* loci is indicated by stars (*). On the right, *MYCN* locus for each tumor is shown with an integrated copy number (CN)/structural variant (SV) plot with absolute copy number on the y-axis and SVs as arcs color-coded by the subclones they were assigned to. *ALK*^ECD-^, *ALK* with a deletion of exons encoding the extra-cellular domain. **b)** Treatment timeline, subclone tree and body map as described in Fig-3a are shown for patient H132384. Evolution at *MDM2, CDK4* and *TERT* loci are shown in the integrated CN/SV plot as described in Fig. 3a. Barplot shows the increase in expression in the tumors with *TERT* SVs. **c)** Treatment timeline, subclone tree and body maps as described in Fig-3a are shown for two different *ATRX*-mutant patients. **d)** Treatment timeline, subclone tree and body maps as described in Fig-3a are shown for two patients with *TP53* mutations. Detailed description of each patient is provided in Supplementary Information. The data for Fig. 3 are available as raw data at dbGAP and scripts are available through ISABL platform.

*TERT*-SVs are mutually exclusive with *MYCN* and *ATRX* rearrangements, thus demarcate a distinct HR-NB subtype^13,17^. In our cohort *TERT*-SVs were enriched in CNS metastases (8/19 patients)^43^. Intriguingly, unlike *MYCN*, where an initial amplification is always present on the trunk, *TERT*-SVs were predominantly (10/13 SVs) subclonal to segmental CNAs (Fig. 2d-e, Fig. 3b and Extended Data Fig. 5). Similar to the *MYCN* locus, we observe continuous *TERT* rearrangements in the majority of cases (5/7) during disease progression. These relapse-specific rearrangements result in increased copy number and expression of *TERT* consistent with an increasing addiction to *TERT* signaling (Fig. 3b, Extended Data Fig. 5 and Supplementary Fig. 3, 5, 18-20 and 24-25).

*ATRX*-mutant neuroblastoma is seen in older patients with indolent disease^44,45^. In contrast to *MYCN-*A and *TERT*-SV patients who were stage-4 at diagnosis, 21% of *ATRX*-mutant cases were diagnosed with low-stage disease but eventually relapsed. Subclonal acquisition of *ATRX* events were common (9/29 events) and seen in relapses (Extended Data Fig. 4). In two patients, parallel acquisition of *ATRX* mutations were seen at distinct metastatic sites (H135089) or locoregional relapses at consecutive time points (H134817) (Fig. 3c). *ATRX*-mutant cases were also enriched for SVs affecting *PTPRD* (Extended Data Fig. 1e) with evidence of parallel or continuous evolution (Fig. 3c and Extended Data Fig. 4).

*MDM2-CDK4* co-amplifications were seen in patients diagnosed with low-stage disease (Supplementary Table 1). Notably, the co-amplification was not mutually exclusive with *ATRX* events and *TERT*-SVs suggesting an overlapping subtype (Fig. 2d-e). For example, phylogenetic reconstruction for patient H132384 mapped truncal *MDM2-CDK4* co-amplification and subclonal SVs at *ATRX* and *TERT*. However, the *ATRX* and *TERT* events were acquired on two distinct subclonal lineages further validating the mutually exclusivity of these events (Fig. 3b). In patients with *MDM2-CDK4*, subclonal *ATRX* events and continuous evolution at *MDM2-CDK4* locus via incorporation and over-expression of other genomic loci (*TERT, WNT3A, IGF2BP3*) were seen during disease progression (Extended Data Fig. 4), which might contribute to the dismal outcomes associated with *MDM2-CDK4*^18^.

*TP53* mutations demarcate an ultra HR-NB subtype^46,47^. Excluding arm-level CNAs at 17p (n=19, 7%), 10 patients had mutations (n=10) or SVs (n=4) at the *TP53* locus specifically. In 7/10 patients these mutations were bi-allelic, most frequently by initial 17p loss followed by a TP53 mutation (n=5). Parallel acquisition of mutations affecting p53 pathway were observed both within the primary site (H116987) and in different metastatic tumors (H134722) (Fig. 3d).

Taken together this analysis demonstrate that in neuroblastoma subtype-specific evolutionary trajectories are predominantly determined by SVs targeting the main driver gene itself with specific acquisition of secondary hits (e.g. *TERTp* in *MYCN-A* and *PTPRD* in ATRX-mutant) and are followed by subclonal mutations in RAS-MAPK, PI3K-mTOR and p53 pathways shared across the disease subtypes.

### Clonal diversification at primary site creates multiple clones with metastatic potential

Patients with HR-NB are diagnosed with widely metastatic disease (bone, bone marrow, liver, lung and CNS)^48^. We studied the clonal relationships amongst primary and disseminated disease in 30 evaluable patients (Methods, Supplementary Tables 3-5).

Whilst all resections were clonally related, subclonal heterogeneity at the primary site was seen in the majority of patients in the form of subclonal CNAs and oncogenic mutations/SVs (83%, 25/30) (CCF median=100%, range=4-100%) (Extended Data Fig. 4-5). This subclonal diversification in the primary site creates distinct cell subpopulations with differential capacity to spread. Analysis of the ensuing metastatic trajectories demonstrates that distinct primary-metastasis pairs share closer lineage relationships in the tumor phylogeny than the primary sites to one another (7/9 patients, Extended Data Fig. 4-5). For example, in patient H103207 (Fig. 3a), clonal structure across two adrenal tumors at diagnosis and 6 metastases from CNS, lungs and liver suggests that CNS-metastatic clone separated from the trunk before the adrenal primary site diversified further. Similarly, one of the two adrenal tumors segregated with the lineage leading to liver and lung metastases. This demonstrates that branching evolution in the primary site gives rise to multiple subclones with the potential to spread. This observation held true across disease subtypes of this cohort (Extended Data Fig. 4-5 and Supplementary Table 3).

### Timing of metastasis with respect to therapy

Detection of therapy-related mutation signatures indicates the presence of cells that survived therapy and subsequently achieved a clonal representation detectable at WGS depth. Therefore, mutational signatures can be used to time emergence of clones relative to the time of therapy^50^. We illustrate this point with patient H118706 (5-yo, *MYCN*-A, stage-4) for whom WGS was performed on two diagnostic tumors (adrenal and liver metastasis) and 15 metastatic sites including liver and lungs at autopsy following unsuccessful treatment with platinum and temozolomide (Fig. 4a). Subclonal structure suggests that all the autopsy tumors came from the same *TP53*-mutant clone with strong exposure to platinum signatures (48% of the substitutions) and this clone was succeeded by six clones with evidence of temozolomide signature (5-19%). This suggests that the *TP53*-mutant clone emerged after platinum therapy and seeded all the metastatic tumors from autopsy when temozolomide therapy started.

**Figure 4.**
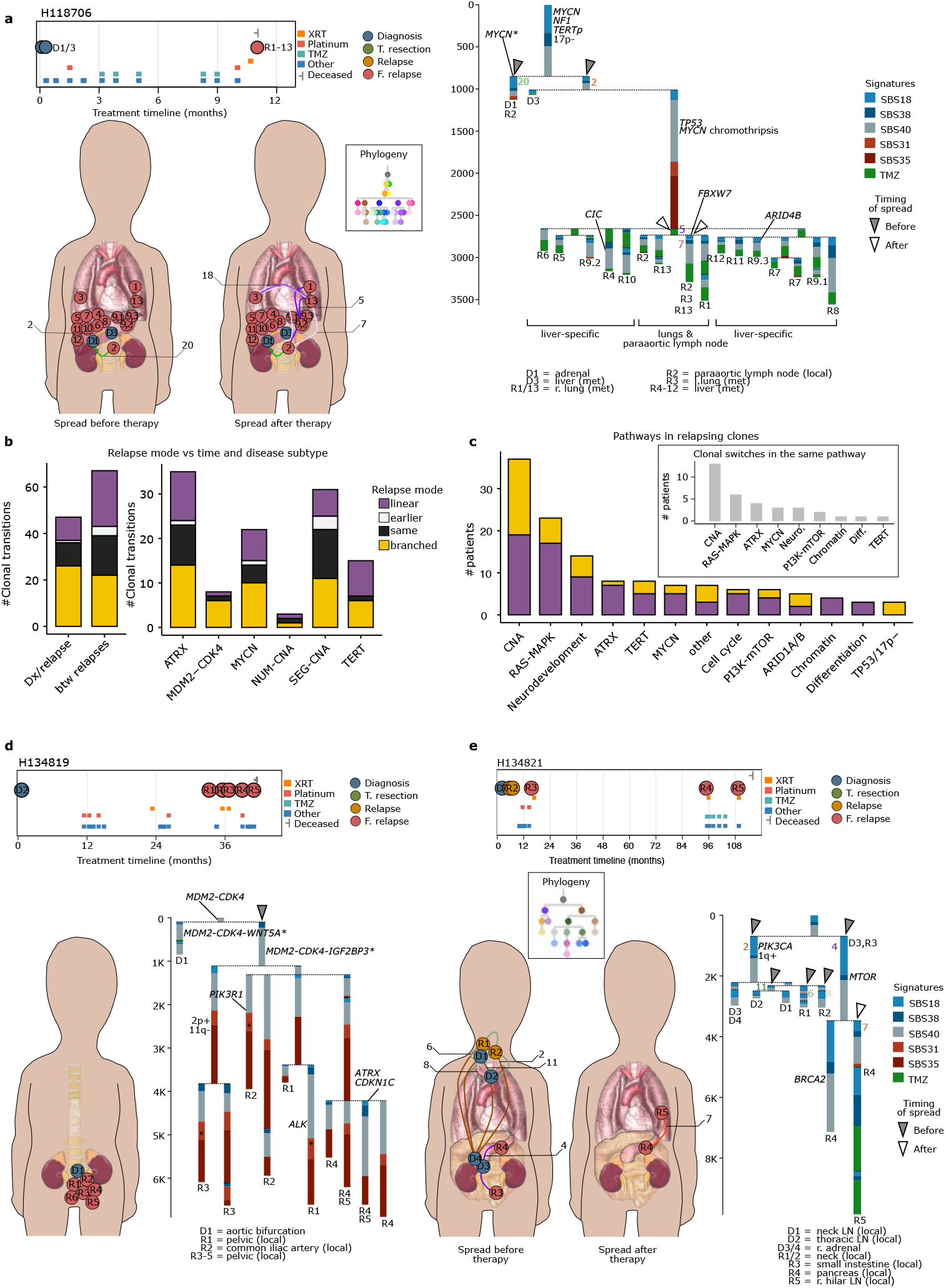
Timing of metastasis. **a)** Subclonal structure for patient H118706 is shown. Treatment timeline is as described in Fig. 3a. On the right is the signature tree with the results from the subclone-specific mutational signature analysis across the subclone tree of the patient. Each subclone is shown as a stacked bar plot showing the proportion of the mutations attributed to the six different mutational signatures and with total length proportional to the number of substitutions in the corresponding subclone. Branches are separated by a dashed line and annotated with the putative oncogenic changes assigned to the corresponding subclone. Terminal nodes are annotated with tumors where the corresponding clone is present. Triangles denote the subclones involved in metastatic spread where gray and white indicate spread before and after therapy, respectively. The id of the metastatic subclones is annotated next to the corresponding branch. Body maps show the possible movement of subclones involved in spread before and after therapy indicated by the subclone ids next to the body maps. Subclones are color-coded according to the clonal phylogeny shown in the legend. Details of the clonal and genomic for H118706 is provided in Supplementary Information. **b)** Left barplot shows the proportion of different types of clonal transitions from diagnosis to first relapse (n=47) and between consecutive relapses (n=67) while the right barplot gives a breakdown of clonal transitions across different disease subtypes. The different types of transitions are 1) ‘same’ where the relapse is caused by the same clone as the previous time point with no new genetic changes acquired. 2) ‘earlier’ where the relapse is caused by an earlier clone in the phylogenetic tree. 3) ‘linear’ where the relapse is caused by a clone with new CNAs and oncogenic mutations/SVs while no such events are seen in the clone specific to the previous time point. 4) ‘branched’ where clones from both time points have genetic changes. **c)** Barplot shows the number of different types of clonal transitions where the relapsing clone has genetic changes affecting the listed CNAs and oncogenic mutations collapsed to pathways affected except for those mutations affecting the most frequent disease-defining genes (*MYCN, TERT* and *ATRX*). The inner plot shows the pathways affected by parallel events across clonal transitions that switch between lineages. **d** and **e)** Subclonal structure for patients H134819 and H134821 are shown. Body maps and treatment timelines are as described in Fig. 3a. Signature trees are as described in Fig. 4a. For H134821 the left body map shows the local-regional spread before therapy while the right body map shows the spread 9 years after diagnosis. Detailed description of each patient is provided in Supplementary Information. The data for Fig. 4a, d and e are available as raw data at dbGAP and scripts are available through ISABL platform while data and scripts for Fig. 4b-c are available in Supplementary Table 6 and the GitHub repository.

Using the same logic, in 13 evaluable patients we studied the relative timing of the emergence of 21 metastasizing subclones and 26 daughter subclones (Supplementary Tables 3-4, Extended Data Fig. 6 and Supplementary Information). Majority of the metastasizing subclones (18/21) had no evidence of therapy exposure, suggesting that disease dissemination happens before therapy consistent with widely metastatic presentation at diagnosis (Extended Data Fig. 6). Importantly, this observation held true not just for local spread but across distant sites including CNS metastases (¾ patients), which are typically not detected by imaging at diagnosis^51^. Notably, exposure to therapy-related mutational signatures was seen in most (22/26) daughter subclones that emerge from the initial metastasizing subclone (CCF median=80%, range=15-100%) in line with on-therapy disease progression at the metastatic sites.

### Origin of late relapses traced back to early clones followed by a long period of dormancy

With improved treatment and prolonged survival, consecutive relapses are increasingly observed amongst HR-NB patients. For 72 patients in our cohort, we were able to study 114 clonal transitions from primary to first relapse (47 patients) and/or between consecutive relapses (43 patients). We observed three distinct patterns of temporal transitions (Supplementary Tables 3 and 6 and Fig. 4b-c). Majority of the relapses were accompanied by accumulation of additional CNAs or mutations/SVs at recurrent loci (72%) (linear and branched in Fig. 4b-c). In 24%, relapses were seeded by exactly the same clone without evidence of new genetic changes and in 4% by an earlier clone in the phylogeny. All three patterns were equally common from diagnosis to first relapse, amongst consecutive relapses as well as across disease subtypes (Fig. 4b). This suggests that while subclones with new drivers continue to emerge and replace existing ones, biological themes are preserved. In 31/45 patients with clonal transitions happening as a switch between different branches, the same pathway was exploited by the tumor consistent with pathway-specific dependencies (Fig. 4c).

For patients with three or more consecutive relapses, we were able to capture multiple waves of clonal successions and transitions across years. For example, for H134819 (3.5-yo stage-I *MYCN*-NA) the same relapsing clone with a *SPRY2* deletion^52^ and a high-level amplification of *IGF2BP3* via *MDM2-CDK4* SVs was present across all five locoregional recurrences from 2.7 to 3.4 years after diagnosis, with late clonal switches from an *ALK* to a *PIK3R1*-mutant subclone and finally to a subclone with *ATRX* and *CDKN1C* events (Fig. 4d). By contrast, in the case of H134821 (9-yo stage-I *MYCN*-NA), the first two locoregional relapses within 4-6 months from diagnosis were caused by a *PIK3CA*-mutant subclone while the last two relapses 8 years later hailed from an *MTOR*-mutant clone (Fig. 4e). Both of these relapsing clones were present at diagnosis suggesting that subclones can stay dormant for many years, remain clinically undetectable but nonetheless maintain the potential to instigate relapses at much later time points. These findings suggest that chemo-resistant clones with specific driver mutations may already exist at diagnosis and, in rare cases, can lay dormant for years before gaining clonal dominance as patients go through multiple lines of therapy.

### Shared lineage relationship across locoregional and distant metastases

We next studied the relationships between locoregional disease and distant metastasis by evaluating data from 19 patients with 69 tumors from disseminated sites (Methods, Supplementary Table 3). We observe an equal number of relapsing subclones involved in locoregional extension or distant metastasis (average 1.5 vs 1.3 subclones) suggesting that there is no differential propensity to extend locally or metastasize. In support of this notion, the same clone from the primary site seeded both locoregionally and at metastatic sites in the four patients with available tumors (30 samples) (Extended Data Fig. 4-5).

### Polyclonal and metastasis-to-metastasis seeding after therapy

Despite high intensity multimodal therapy, 50% of HR-NB patients will eventually progress, typically in metastatic sites. To compare the clonal representation across distant metastatic sites, we studied 10 patients with two or more metastases (Supplementary Table 4). In most cases the same subclone seeded all metastatic tumors (5/7 patients) (Fig. 4a, Fig. 5, Extended Data Fig. 4-5 and Supplementary Information). This occurred in the form of polyclonal seeding^53^ during or after therapy, as evidenced by the presence of therapy related mutation signatures in the metastatic clones (e.g H103207, H118706 and H134722) (Supplementary Information).

**Figure 5.**
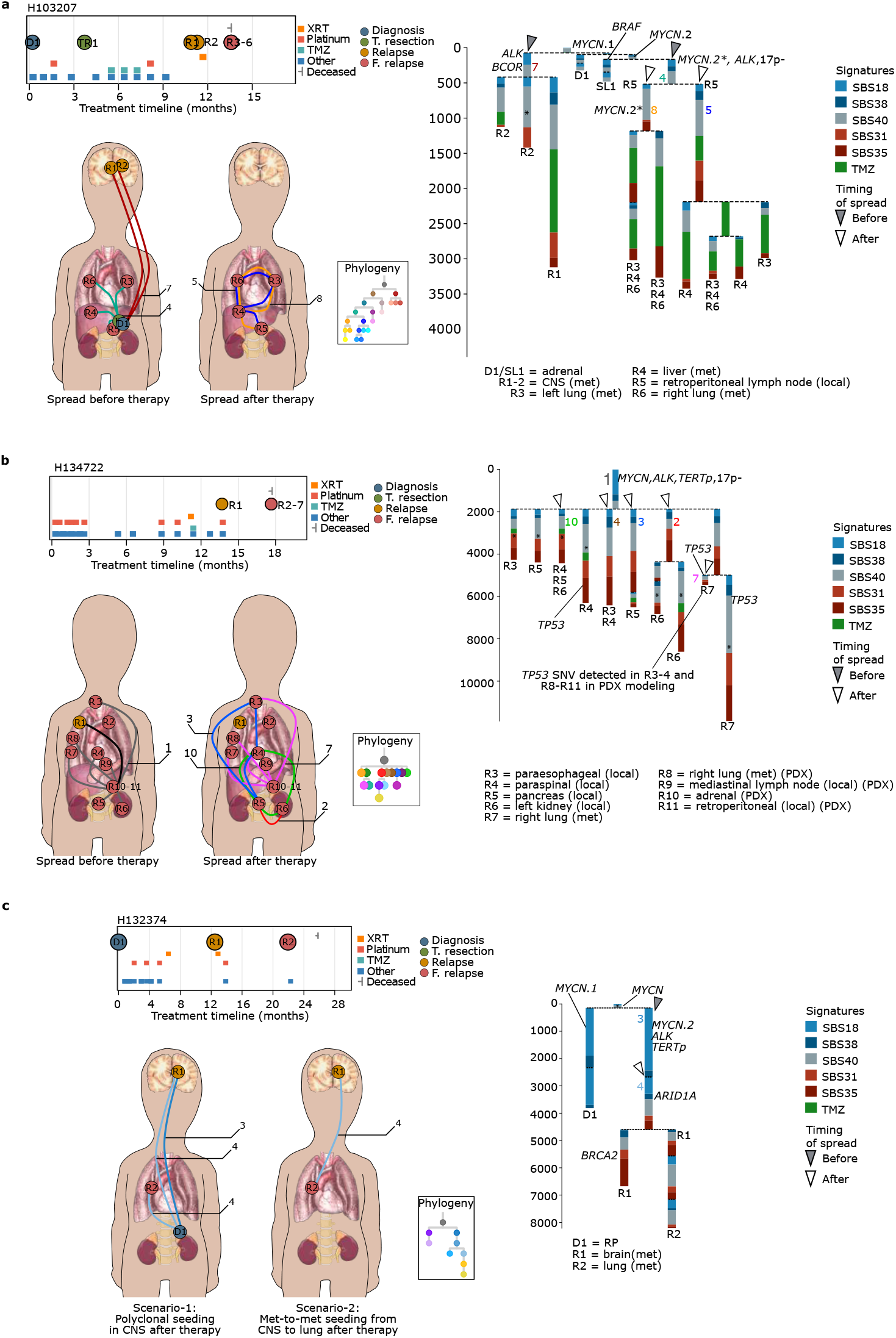
Complex seeding patterns after therapy. Subclonal structures for patients H103207 **(a)**, H134722 **(b)** and H132374 **(c)** are shown with treatment timeline and signature tree as described in Fig. 3a and Fig. 4a, respectively. Detailed description of each patient is provided in Supplementary Information. **a)** Body map on the left shows the seeding events before diagnosis in H103207. Body map on the right depicts the polyclonal seeding after therapy amongst liver and bilateral lungs involving subclones 5 and 8. **b)** The left body map shows the spread before therapy. The black arc indicates the distinct subclone in R1 left lung metastasis sequenced with MSK-IMPACT. The right body map shows subclones 2, 3, 4, 7 and 10 involved in polyclonal seeding across locoregional and metastatic sites in H134722. *TP53* substitution assigned to subclone-7 is also found in PDX modeling of R3, R4, R8, R9, R10 and R11. **c)** Body maps depict two different scenarios that explain the subclonal structure in H132374: The left body map shows possible polyclonal seeding in the CNS by a mixture of subclones 3 and 4. In this scenario lung metastasis is caused by subclone-4 after platinum chemotherapy. Shown on the right body map is the second scenario of met-to-met seeding from CNS to lung by subclone-4 after therapy. The data for Fig. 5 are available as raw data at dbGAP and scripts are available through ISABL platform.

We also see unequivocal evidence of metastasis-to-metastasis (met-to-met) seeding. Patient H118706 was diagnosed with metastatic disease in both the lungs and liver. Interestingly, analysis from 16 tumors from the same metastatic sites 16 months later at autopsy demonstrated that all sites were seeded by the same therapy-exposed subclone originating from the liver metastasis (Fig. 4a). This suggests that the metastatic clones in lungs and liver at diagnosis were cleared by chemotherapy yet a therapy-resistant subclone escaped from the liver and successfully re-seeded multiple metastatic deposits across the same anatomical sites.

Notably the subclones shared across metastases were often characterized by oncogenic events such as the hyper-rearranged *MYCN*-A, convergent *TP53* mutations and *ALK* mutations. This suggests that certain therapy-resistant subclones with selective advantage^54^ have the capacity to disseminate and successfully establish clinically detectable metastatic lesions that transverse locoregional sites, lung, and liver and CNS.

## Discussion

Our understanding of the temporal and spatial relationships underlying evolution of neuroblastoma from disease initiation through treatment, progression and metastasis has remained obscure. Leveraging surgical samples from a unique cohort of 283 patients, we performed a comprehensive analysis of genomic evolution throughout the natural course of the disease.

Clonal reconstruction using multi-sample WGS data traced the first malignant clone and the ensuing subclonal trajectories with notable differences amongst different molecular subtypes. We estimate that MYCN-A emerges during embryogenesis while *ATRX*-mutant disease is frequently post-natal. Unlike *MYCN*, which is always targeted on the trunk, *TERT* and *ATRX* events are not always found in the first malignant clone suggesting that they may not be required for disease initiation but yet, as secondary events, define distinct and non-overlapping clinical subtypes with unique evolutionary trajectories. Once established, tumors progress across time and space via a process underwritten by continuous or parallel acquisition of SVs at disease-defining loci namely *MYCN, TERT, ATRX* and *MDM2-CDK4*. Whilst it is likely that high-level amplifications create genomic instability thereby providing fuel for continued rearrangements, subclonal events at these loci might provide survival advantage under chemotherapeutic pressure, only to become stable at their most complex forms shared across multiple metastatic sites as evidenced by the dominance of clones with the same complex but stable *MYCN* amplicon in late stage metastatic disease. Although this is in agreement with the emerging importance of evolution through extrachromosomal DNA observed in neuroblastoma and other tumor types^56,57^, study of larger sample sets is warranted to further validate these observations.

Analysis of temporally and spatially separated tumors revealed unique patterns and timing of disease spread. At diagnosis, primary disease is already carrying divergent subclones with parallel drivers involved in multiple waves of metastatic spread early in disease evolution. This is in agreement with the idea that clonal heterogeneity is established early in neuroblastoma pathogenesis. As the disease progresses, patients go through multiple lines of therapy, with each consecutive relapse underwritten by the emergence of subclones with new driver mutations affecting the same set of pathways. Furthermore, we show that most metastases occur prior to the commencement of therapy, consistent with the clinical presentation of HR-NB. This is particularly true for CNS metastases which have a low incidence^51^ and are rarely detected clinically at diagnosis^51^. This suggests that CNS involvement becomes detectable only after systemic disease was debulked and controlled by drugs that did not cross the blood brain barrier.

Curative treatment for neuroblastoma is aimed at killing invisible metastases. Indeed we demonstrated that disseminated disease could remain dormant for a long period (up to 10 years) after successful treatment, whilst the patient is clinically “disease-free”, but then eventually relapse and spread to new sites. Despite the observed heterogeneity at the primary site, we show that late metastatic spread after therapy is underwritten by a set of subclones that can spread across locoregional and distant metastatic sites via met-to-met and polyclonal seeding. This suggests that therapy selects for particularly resistant and aggressive subclones with superior metastatic potential.

Taken together our data built a dismal picture of neuroblastoma pathogenesis, where malignant clones arise early in embryogenesis, yet rapidly diversify and spread across local and distant metastatic sites prior to disease diagnosis and unravel the complex networks of disease spread during relapse in response to therapy. This dynamic and rapidly evolving disease presentation has important implications for inclusion of select targeted agents in upfront therapy for HR-NB patients in order to improve the chance of cure^58,59^.

## Online methods

### Patient cohort

Patients in this cohort were seen in the Neuroblastoma clinic at the Memorial Sloan Kettering Cancer Center (MSKCC) from 1987-2021. A written informed consent for tumor/normal sequencing and use of clinical data was taken for all patients in accordance with the ethical rules and regulations of the institutional review board at MSKCC. Additionally three patients’ guardians consented to participate in the MSKCC medical donation program.

### Chemotherapy regimen

Detailed treatment information was available for 56 patients for whom 256 tumors were whole-genome sequenced. Of these, 29 cases received induction chemotherapy as per an MSK protocol (N5, N7, N8 or N9) in which the first five cycles consist of cyclophosphamide/doxorubicin/vincristine (CAV) × 2, cisplatin/etoposide (PVP), CAV, PVP and CAV. 14 patients were treated according to a COG regimen similar to ANBL0532 in which the first five cycles include cyclophosphamide/topotecan × 2, PVP, CAV and PVP while 4 patients were treated according to COG3973 (CAV × 2, PVP, CAV and PVP). The rest of the 8 patients were treated with other protocols containing 3-4 rounds of platinum-based chemotherapy including 1 patient with rapid COJEC. Therapy resection (t-resection) tumors were taken during induction chemotherapy after (3-5) cycles of chemotherapy. Of the 30 patients with WGS data from a t-resection tumor, 21 tumors were exposed to only 1 round of platinum-based chemotherapy while 8 and 1 were exposed to 2 and 3 rounds, respectively.

### Whole genome and transcriptome sequencing data

WGS data for this study cohort came from three different sequencing centers. 1) 45 tumors and matched normals were sequenced at St Judes Children Hospital at a median coverage of 34X for both tumor and normal (range 30-59X)^44^. Publicly available raw data for tumors and normals were downloaded. 2) For 29 tumors and matched normals sequencing library preparation and WGS were performed at the New York Genome Center as described before^60^. Tumors and matched normals were sequenced to a median of 95x (range 73-300X) and 44x (29-88X), respectively. 3) For 173 tumors and matched normals WGS was performed at a median coverage of 83X (40-181X) and 46X (range 36-89X) at MSKCC as described in Supplementary Information. Within the subset of patients where clonal structure was analyzed from multi-WGS data, genome-wide coverage figures were 50-100X for 85% and >100X for 13% of the tumors. Only 6 tumors (all diagnostic) from 6 different patients had <50X coverage. To supplant for lower coverage in the diagnostic tumor at least one additional diagnostic tumor was sequenced whenever available (5/6 patients). RNA sequencing was performed in-house as described in Supplementary Information to achieve a median of 81.5 million paired reads per sample.

### Targeted gene panel sequencing

DNA extracted from formalin fixed paraffin embedded (FFPE) tumor and blood samples (as a matched normal) were sequenced using MSK-IMPACT, an FDA-approved and New York State Department of Health validated panel used to sequence patients’ tumors at MSKCC. MSK-IMPACT captures protein-coding exons of 468 cancer-associated genes, introns of frequently rearranged genes and genome-wide copy number probes^61^. Tumor samples were sequenced at a median depth of 648X, whereas peripheral blood samples at 400x. Established pipelines followed by manual review were used to characterize germline and acquired somatic mutations, copy number variants (CNVs) and if targeted, genomic rearrangements as previously described^61^. Clinically relevant findings were annotated using OncoKb tiers 1-4^62^.

### Bioinformatic analysis of WGS and RNA-seq

Analysis of WGS and RNAseq data was executed using Isabl platform^63^ and included: 1. Data quality control; 2. Ensemble variant calling for germline and somatically acquired mutations from at least two out of three algorithms run for each variant class and 3. Variant classification. Briefly, upon completion of each sequencing run, Isabl imports paired tumor-normal FASTQ files, executes alignment, quality control algorithms and generates tumor purity and ploidy estimates. For tumor samples ensembl variant calling for each variant class (substitutions, insertions and deletions and structural variations) was performed. High confidence somatic mutations are classified with regards to their putative role in cancer pathogenesis and statistical post-processing enables the derivation of microsatellite instability scores and mutation signatures^20^. RNA-seq data were independently analyzed for acquired fusions and gene expression metrics.

Clinical relevance of mutations in common cancer genes was annotated using OncoKb, COSMIC, Ensembl Variant Effect Predictor, VAGrENT, gnomAD and ClinVar databases ^62,64–67^.

Details of the variant calling and annotation can be found in the Supplementary Information.

### Gene expression analysis

Gene expression for 55,390 coding and non-coding genes were ascertained in Transcripts Per Million (TPM) using SALMON (v0.10.0, https://github.com/COMBINE-lab/salmon)^68^. Genes with total expression less than or equal to median across all genes were filtered. Deconvolution of the RNA-seq data to predict the proportion of immune and stromal cells in the tumor microenvironment was done using xCell^72^.

### Identification of mutation signatures for substitutions and indels

De novo mutational signature analysis was performed using sigProfilerExtractor^73^ with default parameters. The signatures identified were compared to the COSMIC Mutational Signatures (v3.2) with the addition of temozolomide signature from Kucab et al^21,74^ using cosine similarity. Details of mutational signature analysis can be found in the Supplementary Information.

### Identification of simple and complex rearrangement events

All structural variants called in a tumor were clustered into simple events (deletion, tandem duplication, unbalanced translocation, balanced translocation, reciprocal inversion) or clustered events (complex with >=2 SVs) using ClusterSV [https://github.com/cancerit/ClusterSV]. The algorithm groups the SVs into clusters based on the proximity of breakpoints, the number of events in the region and the size distribution of those events. The resulting clusters contain SVs that are significantly closer than expected given the orientation and the number of SVs in that tumor and hence are expected to have happened as part of the same event. The resulting clusters were then heuristically refined as described previously^75^. Independently, SV breakpoints and CN data from Battenberg and/or Brass were analyzed using ShatterSeek^76^ to identify chromothripsis events.

### Inference of clonal structure from WGS data

For 45 patients with 170 tumors for whom two or more WGS tumors were available, clonal structure was determined using genome-wide substitutions, indels, SVs and CNAs separately. Within this subset of WGS data, genome-wide coverage figures were 50-100X for 85% and >100X for 13%. Only 6 diagnostic tumors from 6 different patients had <50X coverage. For 5/6 of these patients at least 1 other tumor from diagnosis was sequenced to supplant for lower coverage (Supplementary Information). Phylogenies predicted from substitutions and CNAs are displayed in the Supplementary Figures 2-46.

For substitutions, union of high confidence mutations called across all tumors of the patient was used as input to DPClust (v0.2.2, https://github.com/Wedge-Oxford/dpclust). Mutations were filtered if they 1) had depth greater than 6 standard deviations above median coverage, 2) had no CN information or 3) were in genomic regions that were affected by deletion or copy-neutral LOH in a subset of the tumors of the patient. DPClust algorithm 1) calculates cancer cell fraction (CCF) corrected for purity and local copy number 2) performs clustering across tumors to identify the CCF position for the underlying clusters and 3) assigns mutations to each cluster^77,53^. After heuristic and manual curation of the clusters whenever needed, clusters that predominantly contain mutations located on the chromosome were filtered. Clonal ordering of high-confidence clusters was determined using clonevol (v0.99.11, https://github.com/hdng/clonevol)^78^. When there are multiple possible tumor phylogenies, clones with uncertainty were indicated with a star in the phylogenetic tree. Mutational signatures were computed in each cluster independently. Signature trees were generated with python matplotlib (v3.1.0, https://matplotlib.org/). All steps were run using an inhouse wrapper.

For indels the same filtering criteria as substitutions was used with an additional filtering step prior to clustering. Only indels across loci with tumor depth >=40x and <=200x and with a VAF of >1% were used.

In order to compare the CNA segments in a patient, first aberrant segments are matched to SV breakpoints using the script called “match_rg_patterns_to_library.pl” from the Brass pipeline [https://github.com/cancerit/BRASS/blob/dev/perl/bin/match_rg_patterns_to_library.pl]. The presence of associated SVs across all the tumors (see the next section) is used to determine if CNA breakpoints are shared or tumor-specific. Finally the results are manually cross-checked by comparing the allele-specific and subclonal CN states for the segments as estimated by Battenberg. In addition to this, a separate analysis was performed to construct phylogenies based on CNA segments only using MEDICC2 with default parameters [https://bitbucket.org/schwarzlab/medicc2/src/master/].

### Determining clonal status of SVs in a patient

For all the SVs identified across the tumors of a patient, a pileup procedure was performed to determine the number of aberrant reads supporting the variant in each tumor as described in Supplementary Information. An SV was deemed ‘present’ in tumors with >=2 reads supporting the associated breakpoints. SV clusters were defined as groups of SVs present in the same set of tumors.

### Comparison of substitutions, indel and SV clusters

Substitution and indel clusters were compared by calculating the cosine similarity of the CCF values across all tumors of the patient. Clusters with a cosine similarity >0.9 were matched. When a substitution cluster matched multiple indel clusters, the pair with the smallest summed CCF difference across the tumors was retained. Substitution and SV clusters were compared by their presence across the different tumors of the patient. That is, an SNV cluster present in samples A, B and C is matched with an SV cluster present in the same set of samples.

### Inference of clonal structure from targeted sequencing data

For a subset of 49 patients with 113 tumors that were sequenced with MSK-IMPACT and/or WGS, analysis of clonal structure was confined to the alterations that can be captured by MSK-IMPACT. This includes the substitutions and indels called within the exonic and extended splice site regions of ~450 cancer genes, focal deletions in genes such as *CDKN2A, PTPRD, ATRX* and *TP53*, structural variants in select introns included in targeted sequencing panel in genes such as *ALK, ATRX* and *TP53* as well as arm and chromosome level CNAs. For 29 patients only targeted sequencing data were available while 20 patients had one WGS tumor and at least one tumor sequenced by targeted sequencing. Additional alterations detected by WGS but cannot be captured by MSK-IMPACT (i.e. SVs at loci such as *TERT* and *FOXR1*) are not included in this analysis (indicated with SUBCLONE_TYPE==NA in Supplementary Table 2).

For this patient subset, clonal structure was first analyzed with the allele-specific CNA data as assessed using FACETS^79^ in targeted data and Battenberg in WGS data. CNA-based phylogenies were derived by comparing the genomewide CNAs of the tumors as well as using MEDICC2 with default parameters. Substitutions and indels were analyzed together using DriverClone [https://github.com/papaemmelab/driverclone], an inhouse algorithm designed for studying clonal structure specifically from sparse targeted sequencing data. Briefly, DriverClone first derives a posterior probability for the CCF of each variant, taking into account information on local ploidy, coverage, tumor purity and possible genotypes. DriverClone then clusters the variants in CCF space using a weighted variant graph where edges represent overlaps of posterior credible intervals between variant pairs in each sample. Low weight edges are pruned. A depth-first search then finds all connected components in the variant graph and retrieves clusters of variants belonging to the same predicted clone. To enable probabilistic clonal ordering with few observations per clone, DriverClone extends the non-parametric bootstrapping model of ClonEvol so that bootstrap samples are obtained from a mixture distribution of variant posteriors. A tree enumeration algorithm (originally implemented in Clonevol) then identifies all possible tumor phylogenies that fulfill the appropriate biological constraints. Phylogenies predicted from this analysis are displayed in the Supplementary Figs. 48-53.

### Timing the emergence of MRCA

Timing of the MRCA emergence was performed in a subset of 39 patients with >=2 WGS tumors where at least one tumor was from a pre-treatment diagnostic specimen or therapy resection. Association between age at diagnosis and trunk length was assessed by a linear regression model using R *lm* function and in a multivariate analysis using R *glm* function taking into account disease subtype, stage at diagnosis and number of WGS tumors. MRCA analysis was done using a previously published analysis workflow^37^ in two steps 1) First patient-specific mutation rates were estimated via linear mixed effect modeling with the number of mutations attributed to the clock-like mutational signature SBS40. 2) Patient-specific mutation rates and the number of SBS40 mutations on the trunk were used to estimate the time of emergence for MRCA applying a bootstrapping approach to estimate 95% confidence intervals (CIs). MRCA was classified as ‘pre-natal’ if CIs overlapped the time of birth and ‘post-natal’ otherwise.

### Analysis of truncal and subclonal somatic changes

This analysis was performed in a subset of 94 patients with two or more tumors for which WGS and/or targeted sequencing data was used to assess genome-wide segmental CNAs and oncogenic substitution, indels, CNAs and SVs in reported ~450 cancer genes included in targeted sequencing panel. Detailed analysis of *MYCN* and *TERT* loci was performed in 13 and 7 patients, respectively, with two or more WGS samples available.

### Analysis of evolutionary patterns

Analysis of divergence in the primary site was performed in a subset of 30 patients with two or more tumors from the primary site available. Divergence is defined as acquisition of recurrent CNAs as well as oncogenic mutations provided in Supplementary Table 2. Comparison of primary site to disseminated disease was performed in a subset of 9 patients with two or more tumors from the primary site and tumor(s) from disseminated sites. Timing of metastasis with respect to therapy was performed in a subset of 13 patients for whom at least one tumor from the primary site and two or more tumors from local-regional and/or distant metastatic sites were sequenced by WGS. Lineage relationship between local-regional and/or distant metastatic tumors was studied in a subset of 19 patients with at least one tumor from the primary site as well as local-regional and/or distant metastatic tumors available.

## Author contributions

E.P., N.K.C and G.G. designed the study. G.G., M.F.L., J.S.M.M, J.E.A.O and J.Z. developed algorithmic infrastructure and G.G. performed bioinformatic analysis with support from L.C., M.R. and G.A.. N.K.C, S.S.R., B.S., M.P.L, B.H.K, S.M. and N.S. performed the clinical management of the patients. N.K.C and N.B. performed patient consent. N.K.C oversaw biospecimen banking performed by I.Y.C and Y.F. while D.Y. and F.D.C executed laboratory processing of PDX specimens. N.K.C. collected clinical data for the patients. C.A.I.O. led the clinical donation program. G.G. prepared figures and tables. G.G., N.K.C. and E.P. reviewed analysis results and interpretation of findings and wrote the manuscript with input from D.B.S., B.H.K., S.M., C.A.I.D and A.K.. All authors reviewed and approved the manuscript for submission.

## ACKNOWLEDGEMENTS

The authors would like to acknowledge Drs T. Heaton, and J. Gerstle for their surgical expertise in specimen collections. Dr David Wedge and Dr Maire Ni Leathlobhair of Big Data Institute, University of Oxford, UK, for support and interesting discussions, members of MSK integrative Genomics Operation core for sample processing and sequencing. N-K.C. was partly supported by the Enid Haupt Endowed Chair, the Robert Steel Foundation, Katie Find a Cure, and the Catie Hoch Foundation in building the neuroblastoma tumor tissue archive. E.P. is a Josie Robertson Investigator and is supported by the European Hematology Association, American Society of Hematology, Gabrielle’s Angels Foundation, V Foundation and The Geoffrey Beene Foundation and a Damon-Runyon Rachleff Innovator Award recipient. Funding for this study was supported by the Olayan Fund for Precision Pediatric Cancer Medicine.

## Data availability

All data is available in dbGAP (in submission) and cbioPortal.

## Code availability

Additional scripts and data used for generating the figures are available at https://github.com/gg10/gg10-Clonal-evolution-during-metastatic-spread-in-high-risk-neuroblastoma.

---

**Extended Data Figure 1.**
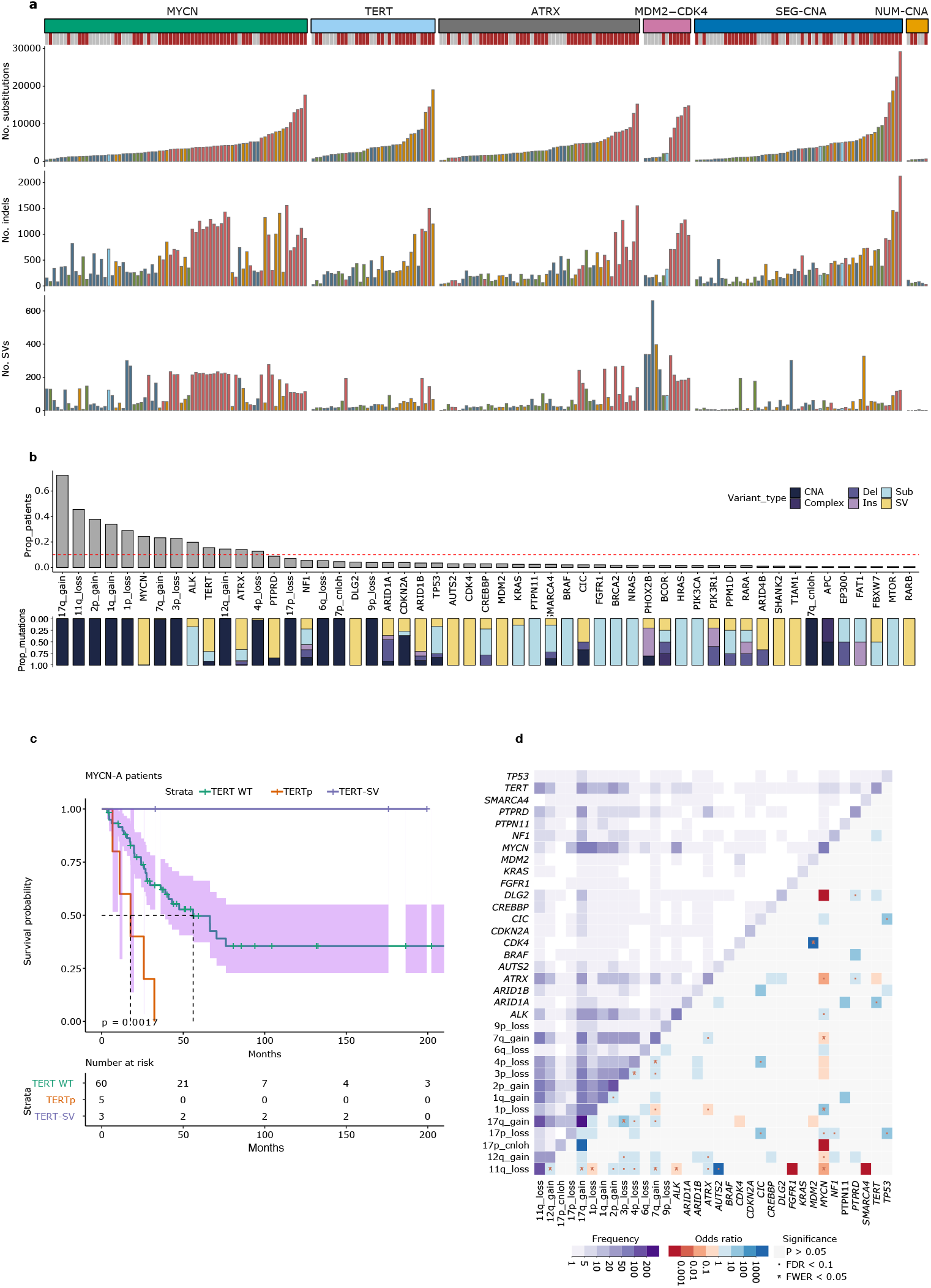
Summary of mutation calling and genomic landscape. **a)** Barplots show the number of substitutions, indels and SVs identified in WGS tumors (n=247) in the cohort grouped by the different disease subtypes and color-coded by the sample type. **b)** Barplots show the prevalence of segmental CNAs and genes affected by mutations and SVs across the cohort (n=470 tumors). Only genes affected in at least two patients are shown. Bottom bar plot gives a summary of the type of mutations for each CNA/gene. CNA, copy number aberration. Complex, small complex insertion/deletion. Del, small deletion. Ins, small insertion. Sub, substitution. SV, structural variant. **c)** Survival plot shows the clinical outcome of *MYCN*-A patients (n=68) with *TERTp* substitutions, *TERT*-SV or no *TERT* events with 95% confidence intervals. P-value from coxph analysis taking into account age at diagnosis is shown. **d)** Heatmap gives a summary of the co-mutation patterns in the current cohort with the frequency of events in the upper triangle and odds ratios in the lower triangle. Only odds ratios with p values <0.05 are colored in shades of blue for co-mutation or red for mutually exclusive interactions. Significant interactions are indicated with a star or a dot according to the significance level. FDR, false discovery rate. FWER, family-wise error rate. The data and script for Extended Fig. 1 are available in Supplementary Tables 1-2 and the GitHub repository.

**Extended Data Figure 2.**
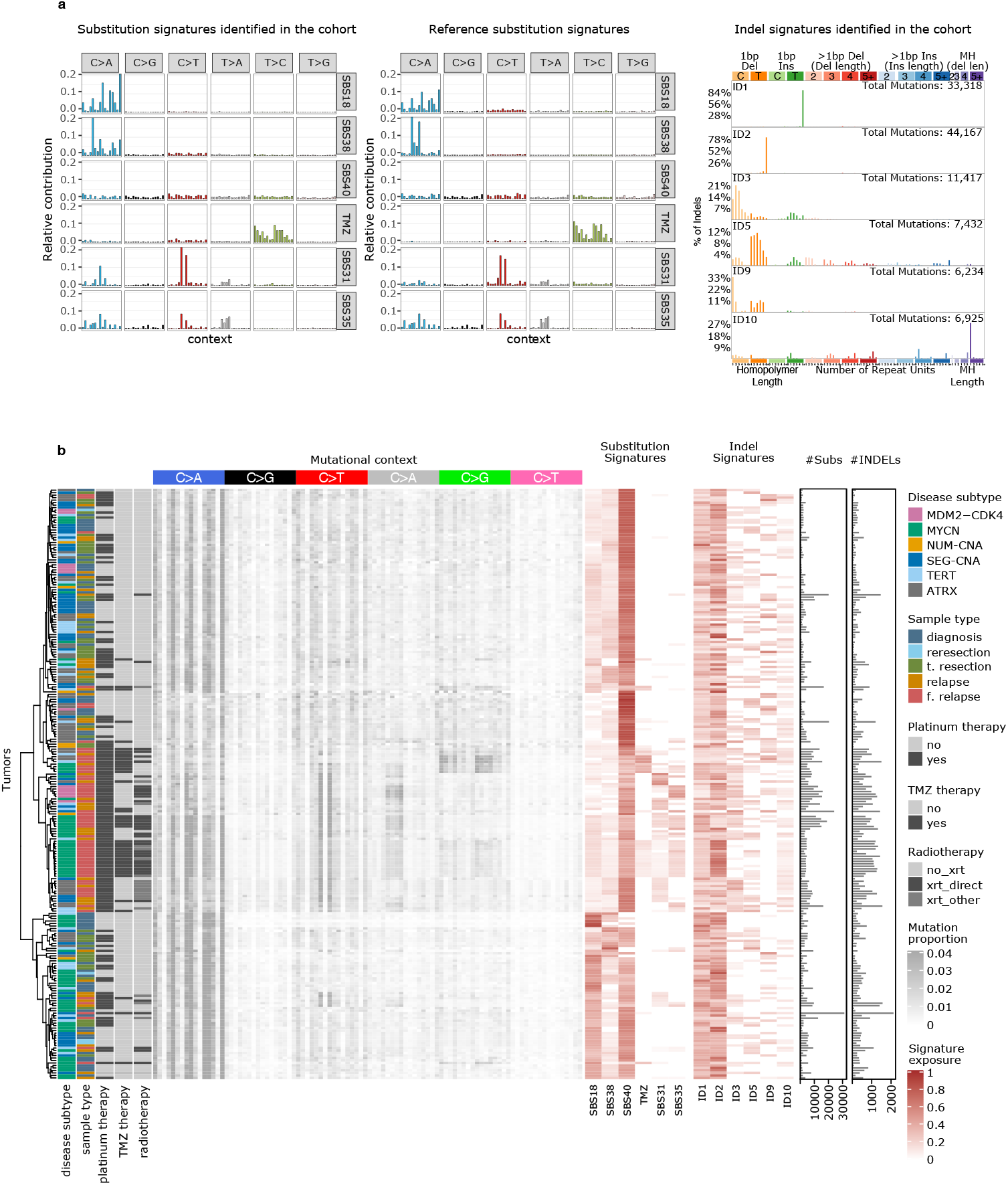
Summary of mutational signature analyses in the current WGS data. **a)** 96 mutational contexts for the substitution signatures identified de novo are shown as the barplots on the left while the reference signatures from COSMIC.v3 are shown in the barplot in the middle. Right barplot shows the prevalence of different types of indels amongst the indel signatures identified de novo. **b)** Heatmap shows the proportions of substitutions at 96 mutational contexts for each WGS tumor (n=247) shown in rows together with sample type, disease subtype, platinum, temozolomide and radiotherapy status on the left and number of substitutions and indels and exposure to identified signatures in substitution and indel data on the right. The data and script for Extended Fig. 2 are available at the GitHub repository.

**Extended Data Figure 3.**
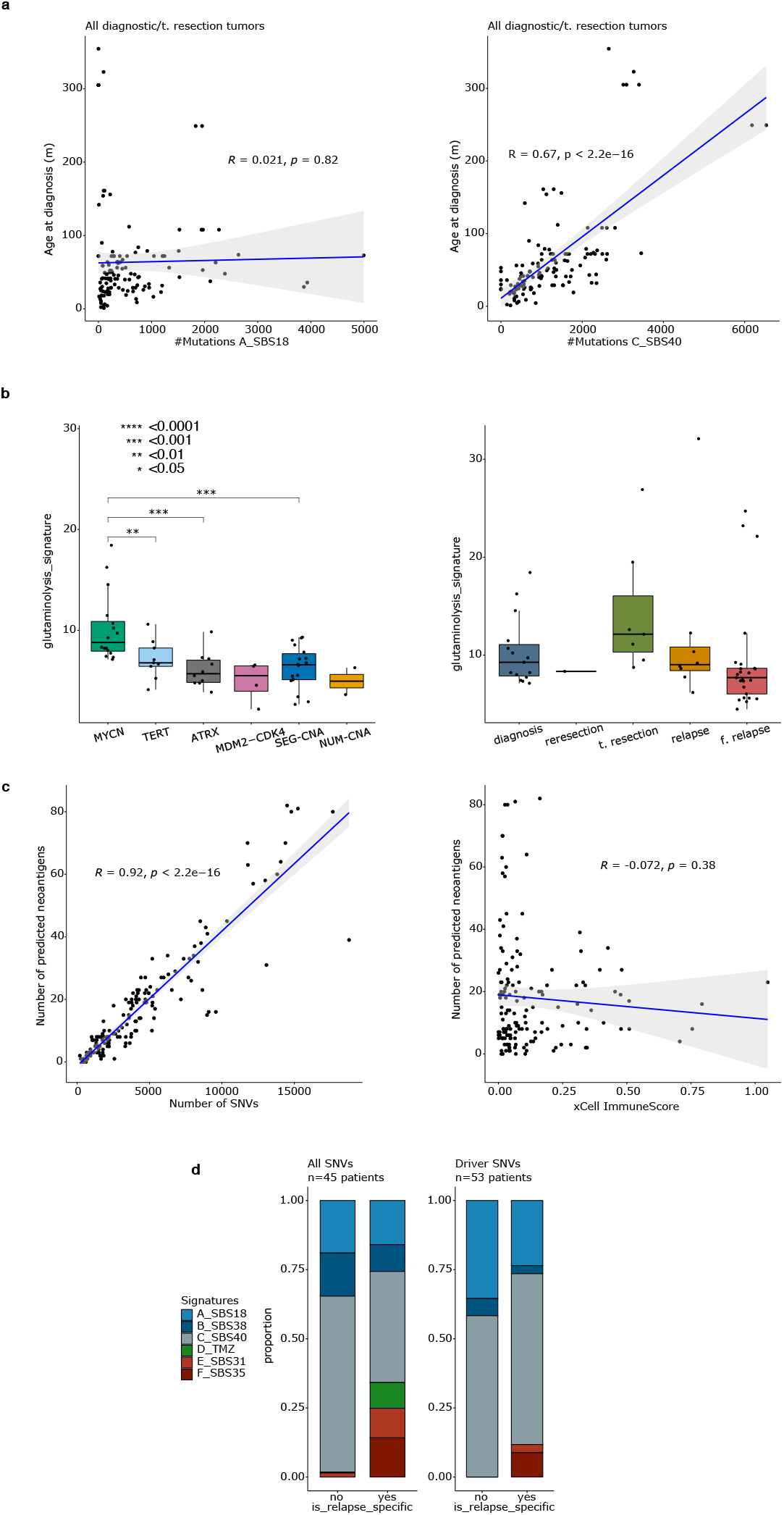
Biological and clinical correlates of mutational patterns. **a)** Scatter plots show the association between exposure to SBS18 (left) or SBS40 (right) and age at diagnosis amongst the diagnostic/t-resection tumors (Pearson correlation). **b)** Box plot shows the mean expression of the genes in glutaminolysis signature associated with ROS accumulation^22^ across diagnostic tumors of different disease subtypes (left) and *MYCN-A* tumors from diagnosis, t-resection and relapse and further relapses (right). Comparisons with significant p-values are shown. **c)** Scatter plot on the left shows correlation between number of SNVs and the number of predicted neoantigens while the scatter plot on the right shows the relationship between the number of predicted neoantigens and immune infiltrates in the surrounding tumor microenvironment as assessed from RNAseq (Pearson correlation). **d)** Barplots show the proportion of genome-wide SNVs (left) and oncogenic driver SNVs (right) attributed to different mutational signatures broken down by presence in post-therapy relapse tumors. The data and script for Extended Fig. 3 are available in Supplementary Table 1 and the GitHub repository.

**Extended Data Figure 4-5.**
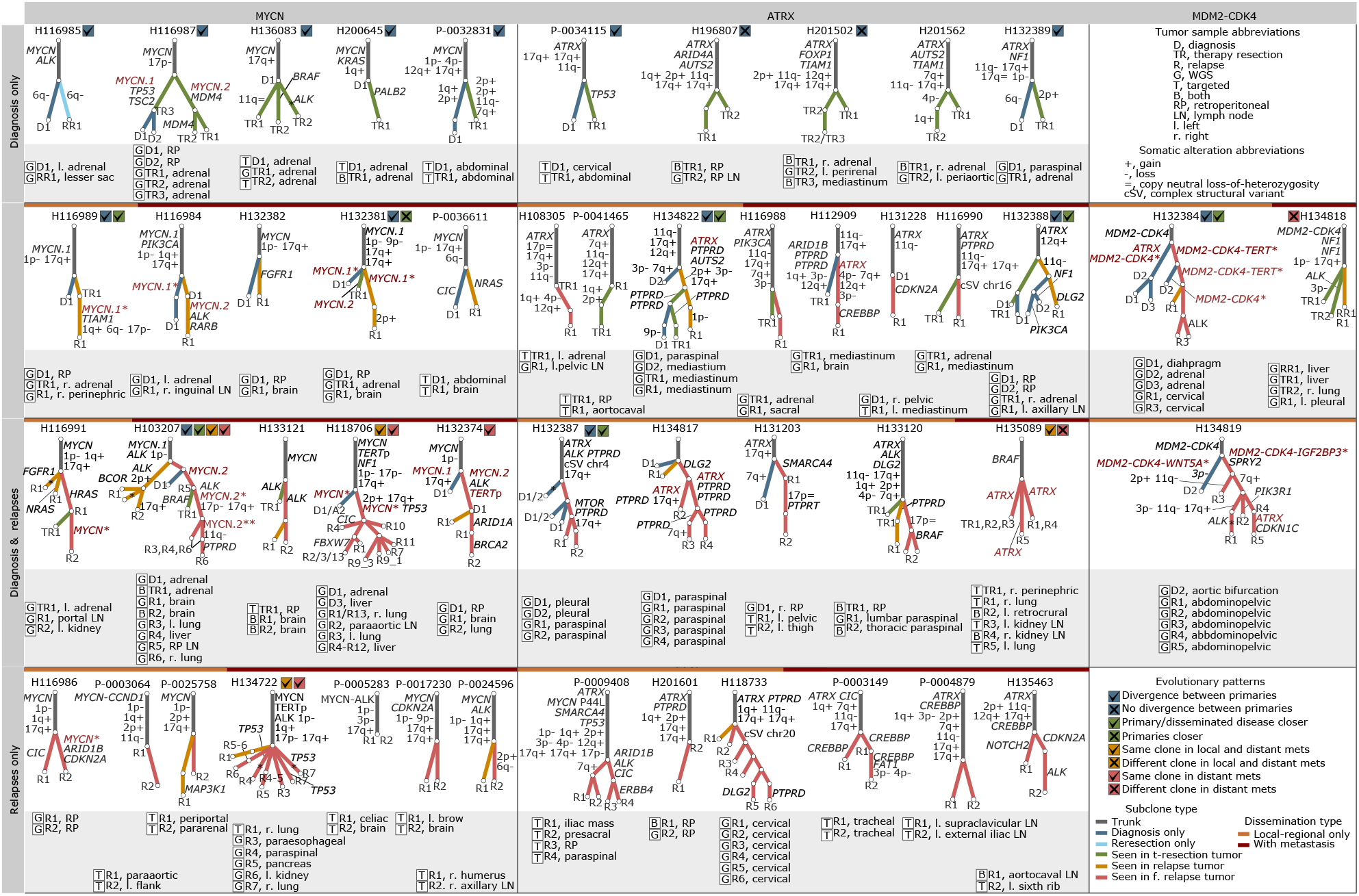
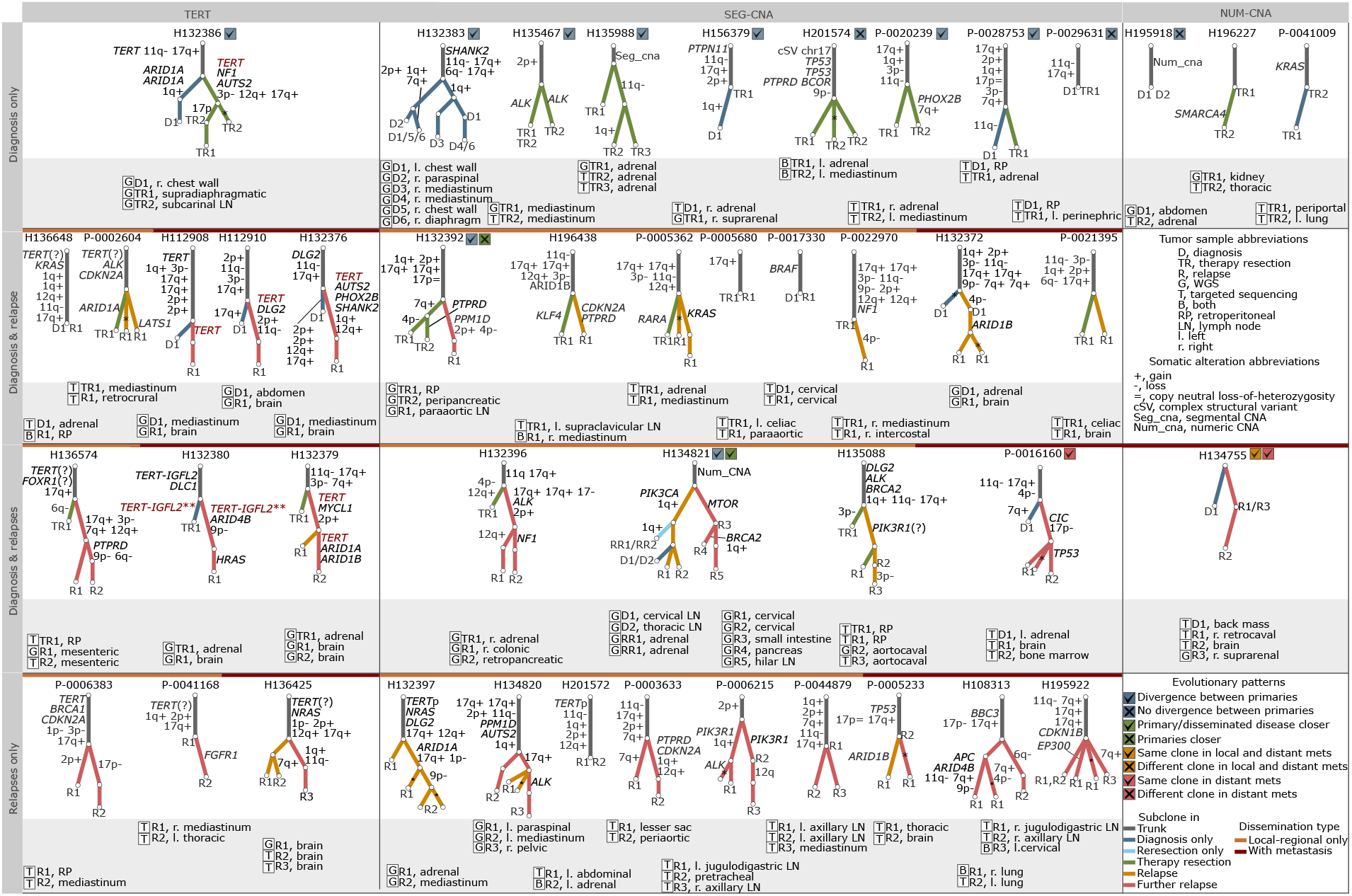
Subclone trees for 94 neuroblastoma patients with two or more tumors sequenced. Each tree shows the subclonal structure in an individual patient. Patients are organized according to disease subtype and the availability of tumors from primary site (diagnosis/reresection/t-resection) and relapses. Branches are annotated with recurrent CNAs and oncogenic mutations/SVs and colored according to the latest tumor they were identified in 1) blue for subclones specific to a diagnostic tumor 2) light blue for subclones seen in reresections 3) green for subclones seen in a t-resection tumor 4) orange for subclones seen in a relapse tumor and 5) red for subclones seen in a further relapse tumor. Subclonal events at *MYCN, TERT* and *ATRX* loci are shown in red font. Events with which clonal status cannot be determined are indicated with a question mark. Different evolutionary patterns are indicated with an icon next to the patient id. Tumor sites and the type of sequencing are indicated below the trees. G, whole-genome sequencing. T, targeted sequencing. B, both. For H103207, H118706 and H134819 a simplified version of the tree is shown due to space. Detailed analysis of subclonal structure for 94 patients is provided in Supplementary Fig. 2-46 and 48-53. The data Extended Figs. 4-5 are available in Supplementary Table 5 and the scripts are available through the ISABL platform.

**Extended Data Figure 6.**
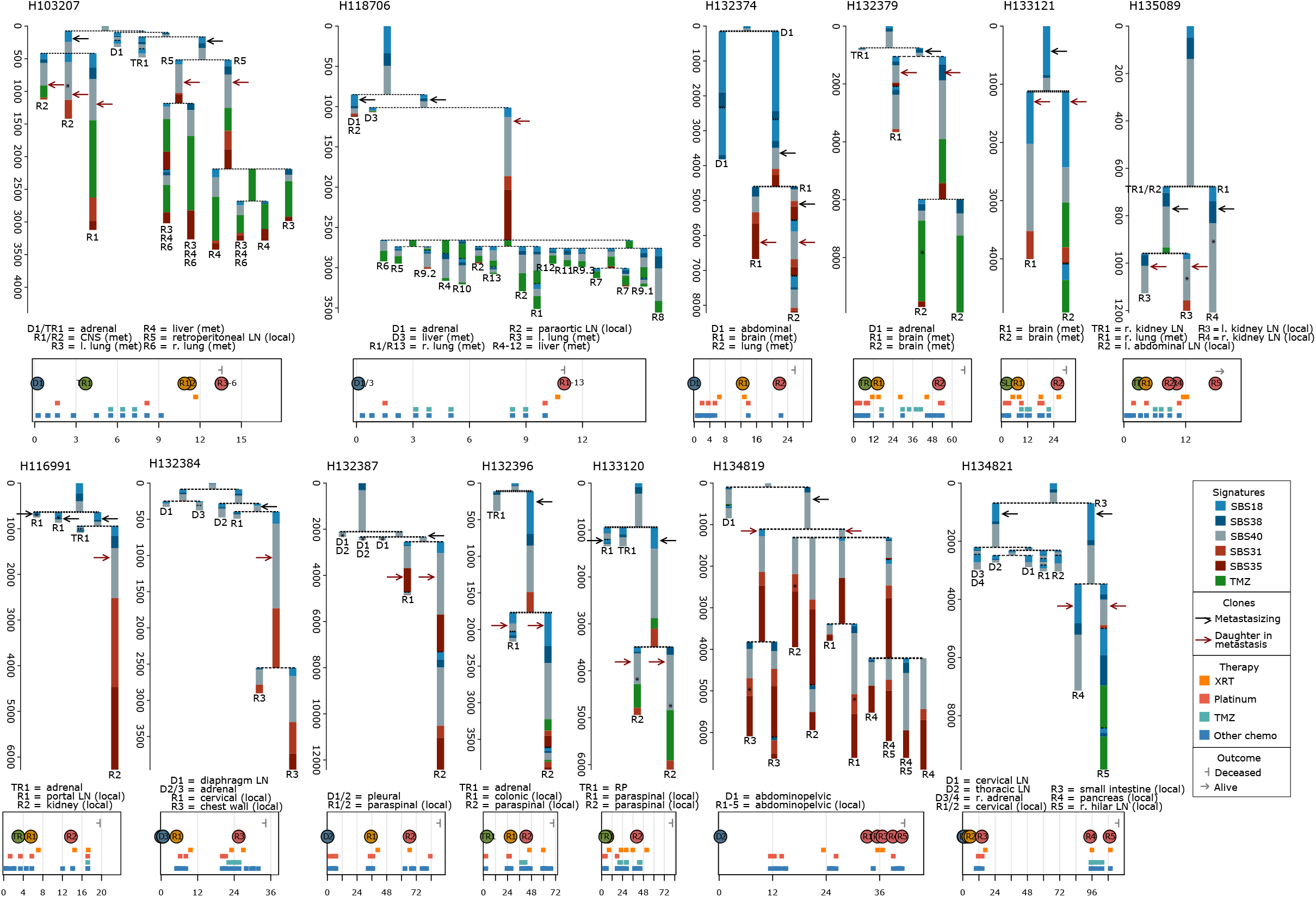
Timing of metastasis. Signatures trees as described in Fig-4a are shown for 13 patients with one or more tumors from the primary site and two or more tumors from locoregional and/or distant metastasis. Patients in the top row have at least one tumor from distant metastatic site while patients in the bottom row have locoregional relapses only. The subclones involved in disease spread from the primary are indicated with a black arrow while the daughter clones are shown with red arrows. The data for Extended Fig. 6 are available as raw data at dbGAP and the scripts are available through the ISABL platform.

